# Loci associated with cave-derived traits concentrate in specific regions of the Mexican cavefish genome

**DOI:** 10.1101/2024.03.29.587360

**Authors:** Jonathan Wiese, Emilie Richards, Johanna E. Kowalko, Suzanne E. McGaugh

**Affiliations:** Department of Ecology, Evolution, and Behavior, University of Minnesota, St. Paul, MN; Department of Biological Sciences, Lehigh University, Bethlehem, PA

## Abstract

A major goal of modern evolutionary biology is connecting phenotypic evolution with its underlying genetic basis. The Mexican cavefish (*Astyanax mexicanus*), a characin fish species comprised of a surface ecotype and a cave-derived ecotype, is well suited as a model to study the genetic mechanisms underlying adaptation to extreme environments. Here we map 206 previously published quantitative trait loci (QTL) for cave-derived traits in *A. mexicanus* to the newest version of the surface fish genome assembly, AstMex3. This analysis revealed that QTL cluster in the genome more than expected by chance, and this clustering is not explained by the distribution of genes in the genome. To investigate whether certain characteristics of the genome facilitate phenotypic evolution, we tested whether genomic characteristics, such as highly mutagenic CpG sites, are reliable predictors of the sites of trait evolution but did not find any significant trends. Finally, we combined the QTL map with previously collected expression and selection data to identify a list of 36 candidate genes that may underlie the repeated evolution of cave phenotypes, including *rgrb* which is predicted to be involved in phototransduction. We found this gene has disrupted exons in all non-hybrid cave populations but intact reading frames in surface fish. Overall, our results suggest specific “evolutionary hotspots” in the genome may play significant roles in driving adaptation to the cave environment in *Astyanax mexicanus* and demonstrate how this compiled dataset can facilitate our understanding of the genetic basis of repeated evolution in the Mexican cavefish.

## Introduction

A major goal of modern evolutionary biology is connecting phenotypic evolution with its underlying genetic basis. Understanding the genetic basis of trait evolution can help reveal whether shared or different genetic mechanisms contribute to repeated trait evolution (Arendt and Reznick 2008; Martin and Orgogozo 2013; Manceau et al. 2010; Waters and McCulloch 2021) and how often repeated evolution occurs through the same or divergent genetic variants (Courtier-Orgogozo et al. 2020; Lee and Coop 2019, 2017). Further, while studies have revealed that loci contributing to trait evolution can cluster in specific genomic regions, the extent to which specific regions in the genome are commonly used in trait evolution and what features of these regions make them more likely to harbor genetic variants contributing to phenotypic evolution remains understudied (Woodhouse and Hufford 2019; Storz 2016; Wortel et al. 2023; Xie et al. 2019).

Linking genotypes to specific phenotypes remains the first step in any query of the repeatability of molecular evolution, and quantitative trait locus (QTL) mapping is a powerful technique for discovering statistical associations between genotypes and phenotypes that has been utilized since the 1980’s (Paterson et al. 1988) on a wide variety of organisms and traits (Yano et al. 1997; Nuzhdin et al. 2005; Shaw et al. 2007). QTL mapping is often the first step to understanding proximate mechanisms for how traits evolve, however, any single QTL study will underestimate the number of loci that influence a given trait and often overestimate their effect sizes (Beavis 1994; Kearsey and Farquhar 1998; Xu 2003). Combining the results of several QTL studies for the same trait can identify loci that were missed by a single study and give a more comprehensive estimate of the number of loci underlying the trait (Peichel and Marques 2017).

The Mexican cavefish, *Astyanax mexicanus*, is an ideal model system to investigate questions regarding the genetic basis of repeated adaptation to novel environments. *Astyanax mexicanus* is a species of freshwater characin fish that exists in two distinct ecotypes: a surface form found in streams across southern Texas and northeastern Mexico, and a cave-dwelling form comprised of populations in at least 35 different caves (Espinasa et al. 2020; Elliott 2018; Miranda-Gamboa et al. 2023). Phylogeographic analyses suggest that cave populations arose from at least two distinct colonization events from two distinct surface ancestral lineages within the last 200,000 years, and that since the initial colonization most cave populations have evolved relatively independently from one another, although there is evidence of gene flow between some cave populations (Moran et al. 2023; Herman et al. 2018; Coghill et al. 2014; Bradic et al. 2012). To varying degrees, cave populations have convergently evolved common phenotypes consisting of morphological traits, most notably degenerated eyes and reduced pigmentation (Jeffery 2001, 2020), as well as behavioral characteristics such as decreased sleep, schooling, and stress behaviors (Kowalko et al. 2013b; Duboue et al. 2011; Chin et al. 2019). The most frequently studied cave populations include the Pachón and Tinaja cave populations, originated from caves located in the Sierra de El Abra region of Mexico, and the Molino cave population, from the nearby Sierra de la Guatemala (Jeffery 2020). Importantly, fish from cave-dwelling *A. mexicanus* populations can interbreed with fish from surface populations (Jeffery 2001; Jeffery 2008). The ability to cross cave and surface-dwelling fish to generate viable hybrid offspring allows researchers to utilize QTL mapping to identify statistical associations between genotypes and phenotypes, facilitating the study of the genetic basis of phenotypic evolution in *A. mexicanus* (Jeffery 2001; Jeffery 2008; O’Quin and McGaugh 2015).

In the last two decades, 20 QTL studies have been conducted in efforts to elucidate the genetic bases of traits in *A. mexicanus* (Table 1). The first QTL study in *A. mexicanus* was published by Borowsky and Wilkens (2002) who identified QTL for three quantitative traits: eye size, pigmentation, and condition factor (a measure of weight scaled by length). Intriguingly, this study identified two clusters of QTL, one consisting of overlapping QTL for condition factor and pigmentation and another consisting of overlapping QTL for condition factor and eye size. In their discussion, Borowsky and Wilkens (2002) highlighted the overlapping QTL and raise two possible explanations: (1) that the same genes underlie both traits due to pleiotropic effects, or (2) that different closely linked genes underlie the traits. A series of QTL studies by Protas and colleagues (Protas et al. 2008; Protas et al. 2007; Protas et al. 2006) corroborated the findings of Borowsky and Wilkens and showed that many QTL for cave-derived traits form clusters in the *A. mexicanus* genome. Since these initial studies, (O’Quin and McGaugh 2015) mapped all published QTL (through 2014) to the first draft of the *A. mexicanus* Pachón cavefish genome (McGaugh et al. 2014), and found 13 significant QTL clusters across eight linkage groups.

**Table 1.** Summary of published Quantitative Trait Loci mapping studies in *Astyanax mexicanus*. SF = surface fish. BC = backcross (F1 hybrids x Parental generation), F2 = second filial generation (F1 hybrids x F1 hybrids), F3 = third filial generation (F2 hybrids x F2 hybrids). In some cases, the number of pigmentation QTL that are mapped may be inflated because pigmentation at different places on the body were mapped as separate traits and the QTL overlapped, yet, we count them separately for this table. 1. Protas et al. (2006) and Gross et al. (2009) used the same Molino x SF backcross (n=111) 2. Protas et al. (2006), Protas et al. (2007), Protas et al. (2008), Gross et al. (2009), Gross et al. (2014), and Lyon et al. (2017) used the same Pachón x SF F2 mapping population (n=539) 3. Yoshizawa et al. (2012), O’Quin et al. (2013), O’Quin et al. (2015), Yoshizawa et al. (2015), and Lyon et al. (2017) used the same Pachón x SF F2 mapping population (n=384) 4. Carlson et al. (2015), Carlson et al. (2018), and Powers et al. (2020) used the same Pachón x SF F2 mapping population (n=129). 5. Riddle et al. (2020), Riddle et al. (2021), and Warren et al. (2021) used the same Pachón x SF F2 mapping population (n=219). 6. Stockdale et al. (2018) and Warren et al. (2021) used the same Pachón x SF F2 mapping population (n=188)

| Study | Populations | Cross | Trait | # of QTL identified |
| --- | --- | --- | --- | --- |
| Borowsky and Wilkens (2002) | Pachón x SF | BC | Albinism | 1 |
|  |  |  | Condition Factor | 2 |
|  |  |  | Eye Size | 3 |
|  |  |  | Pigmentation | 2 |
| Protas et al. (2006) | Molino x SF<sup>1</sup>, Pachón x SF<sup>2</sup> | BC, F2 | Albinism | 1 |
| Protas et al. (2007) | Pachón x SF<sup>2</sup> | F2 | Eye Size | 8 |
|  |  |  | Jaw bone length | 7 |
|  |  |  | Lens size | 6 |
|  |  |  | Maxillary tooth number | 6 |
|  |  |  | Pigmentation | 18 |
|  |  |  | Taste bud number | 6 |
| Protas et al. (2008) | Pachón x SF<sup>2</sup> | F2 | Eye size | 8 |
|  |  |  | Pigmentation | 4 |
|  |  |  | Condition factor | 9 |
|  |  |  | Weight loss | 1 |
|  |  |  | Tooth number | 5 |
|  |  |  | Peduncle depth | 3 |
|  |  |  | Fin placement | 9 |
|  |  |  | Anal fin rays | 2 |
|  |  |  | SO3 width | 6 |
|  |  |  | Rib number | 6 |
|  |  |  | Length | 5 |
|  |  |  | Chemical sense | 4 |
| Gross et al. (2009) | Molino x SF<sup>1</sup>, Pachón x SF<sup>2</sup> | BC, F2 | Brown melanophore | 1 |
| Yoshizawa et al. (2012) | Pachón x SF<sup>3</sup> | F2, F3 | Albinism | 1 |
|  |  |  | Eye size | 5 |
|  |  |  | Vibration attraction | 2 |
|  |  |  | Superficial neuromasts | 2 |
| O'Quin et al. (2013) | Pachón x SF<sup>3</sup> | F2 | Retinal thickness | 4 |
| Kowalko et al. (2013a) | Pachón x SF, Tinaja x SF | F2 | Feeding angle | 2 |
|  |  |  | Jaw angle | 1 |
|  |  |  | Eye size | 5 |
|  |  |  | Head depth | 1 |
|  |  |  | Feeding angle | 1 |
|  |  |  | Tastebuds | 2 |
|  |  |  | Body depth | 1 |
|  |  |  | Eye orbit diameter | 5 |
| Kowalko et al. (2013b) | Tinaja x SF | F2 | Schooling behavior | 2 |
|  |  |  | Dark preference | 1 |
|  |  |  | Eye size | 1 |
|  |  |  | Pupil size | 1 |
| Gross et al. (2014) | Pachón x SF<sup>2</sup> | F2 | Albinism | 1 |
|  |  |  | Eye size | 2 |
|  |  |  | Craniofacial bones | 12 |
|  |  |  | Sex | 1 |
|  |  |  | Standard length | 1 |
| Carlson et al. (2015) | Pachón x SF<sup>4</sup> | F2 | Albinism | 1 |
| O'Quin et al. (2015) | Pachón x SF<sup>3</sup> | F2 | Scleral ossification | 1 |
|  |  |  | Scleral width | 1 |
|  |  |  | Eye size | 8 |
| Yoshizawa et al. (2015) | Pachón x SF<sup>3</sup> | F2, F3 | Vibration attraction | 2 |
|  |  |  | Locomotor activity | 2 |
| Lyon et al. (2017) | Pachón x SF<sup>2,3</sup> | F2 | Scleral ossification | 3 |
| Carlson et al. (2018) | Pachón x SF<sup>4</sup> | F2 | Velocity | 8 |
|  |  |  | Top usage | 2 |
|  |  |  | Bottom usage | 5 |
| Stockdale et al. (2018) | Pachón x SF<sup>6</sup> | F2 | Heart regeneration | 3 |
| Powers et al. (2020) | Pachón x SF<sup>4</sup> | F2 | Eye orbit neuromasts | 1 |
|  |  |  | Eye orbit area | 1 |
| Riddle et al. (2020) | Pachón x SF<sup>5</sup> | F2 | Carotenoid accumulation | 1 |
| Riddle et al. (2021) | Pachón x SF<sup>5</sup> | F2 | Blood glucose | 2 |
|  |  |  | Weight | 7 |
|  |  |  | Length | 5 |
|  |  |  | Condition factor | 3 |
|  |  |  | Gonad weight | 9 |
|  |  |  | Liver area | 6 |
|  |  |  | Gut length | 5 |
| Warren et al. (2021) | Pachón x SF<sup>5,6</sup> | F2 | Albinism | 2 |
|  |  |  | Eye size | 2 |

Since this 2015 analysis (O’Quin and McGaugh 2015), the number of published QTL studies for *A. mexicanus* has nearly doubled, with dozens of new QTL for traits ranging from scleral ossification to heart regeneration (Table 1). Further, a new surface fish chromosome level genome assembly, AstMex3, was made available in July 2022. Mapping QTL to a more complete surface fish genome, instead of the cavefish genome, will yield more comprehensive results if important loci were missing from the fragmented cavefish assembly. The first draft cavefish genome was a scaffold level assembly, comprised of 10,735 scaffolds with a contig N50 of 14.7 kb and a scaffold N50 of 1.775 Mb (McGaugh et al. 2014). In contrast, the AstMex3 surface fish assembly is chromosome level, containing 25 chromosomes, 109 unplaced scaffolds, and greatly improved N50 values of 47.1 Mb for contigs and 51.9 Mb for scaffolds (Warren et al. 2021). Thus, linkage between traits as well as additional candidate genes may be uncovered since the new genome is much more contiguous.

Here we present an updated analysis of the distribution of QTL across the *A. mexicanus* genome, incorporating most published QTL studies in this species to date. Genetic markers used in prior studies were re-mapped to the AstMex3 genome to create an integrated genomic map, which was utilized to anchor QTL intervals to the new genome. Putative QTL clusters were then identified. To understand if certain mutational biases predispose regions to contribute to phenotypic evolution, we tested several genomic variables for correlations with the observed QTL. Next, while acknowledging the bias of QTL to identify regions of moderate to large effect (Rockman 2012), we analyzed the percent variance explained (PVE) values of cave-derived QTL for different phenotypic classes. Lastly, we demonstrate the utility of our compiled QTL analysis for the evolutionary genetics community by cross-referencing QTL for cave-derived phenotypes with other sources of genomic data to identify candidate genes for repeated trait evolution in cavefish. In sum, this study contributes to our understanding of adaptation to an extreme environment and provides a resource for expedited candidate gene discovery and understanding potential links between multiple phenotypes.

## Methods

### Literature review

Quantitative trait locus (QTL) mapping studies in *Astyanax mexicanus* were identified by querying the online databases PubMed and Web of Science with the following search terms: “astyanax” AND “qtl”; “cavefish” AND “qtl”; “astyanax” AND “quantitative trait loc*”; “cavefish” AND “quantitative trait loc*”. The resulting publications were manually filtered to exclude studies that did not map de novo QTL, as well as studies for which genetic marker sequences were not available. Only studies published before 6/6/22 were included in the analysis.

### Mapping QTL to the AstMex3 genome

To map previously published QTL to the most recent version of the *A. mexicanus* genome, AstMex3, all available marker sequences used in prior QTL studies were collected and aligned to the AstMex3 surface genome with BLAST+ version 2.8.1 (Camacho et al. 2009) (Table S1, S2, S3). For each marker sequence, the best quality BLAST alignment was identified by filtering according to the following criteria: 1. Highest bitscore, 2. Smallest e-value, 3. Highest percent identity. We refer to this match as the “best hit.” Markers with multiple BLAST hits that could not be ranked using these criteria were discarded from the analysis. Additionally, markers for which the best BLAST hit mapped to a chromosome discordant with other markers on its original linkage group were discarded. The best BLAST hits for each of the remaining markers were collated into an integrated genomic map, anchored to the AstMex3 surface genome (Table S4). Three genes (*igf1*, *shhb* also known as *twhh, pax6*) that were used as markers in previous QTL studies but did not have sequences included in our marker database were added manually to the integrated map using the known genomic coordinates given in the AstMex3 genome assembly.

The integrated genomic map was used to identify the location of each QTL in the AstMex3 surface genome assembly. We caution that this is a rough estimate of QTL location, as we were comparing across over 20 years of studies, many of which report different levels of information. Nevertheless, compiling QTL peaks, even if rough approximation across studies, may be useful for future candidate gene identification and to understand traits with potentially co-localizing loci. When available, the genetic markers designating the 95% confidence interval for each QTL (as defined by the study in which the QTL was originally mapped) were used to designate the section of the genome in which loci underlying the trait of interest may be found (see Table S5 for available metadata for each QTL). If a confidence interval was not available, only the marker representing the QTL peak was used to estimate the location of the QTL in the genome. If the markers designating both the 95% confidence interval and the peak of a given QTL were absent from the integrated genomic map, then that QTL was discarded from the rest of the analysis. We manually assigned the phenotypic category of each QTL that was mapped to the AstMex3 genome into one of eight categories (Eye, Pigment, Skeletal, Non-Eye Sensory, Metabolic, Behavior, Cardiac, Other morphological (this category consisted of traits that did not fit clearly into one of the other seven categories, e.g. length); Tables S5, S6), and the QTL intervals and peaks mapped to each chromosome were visualized using the karyoploteR package (v. 1.24.0; Gel and Serra 2017).

### Identifying QTL clusters

To analyze QTL distribution across the genome, QTL were filtered to include only the QTL that were mapped to intervals (not just peaks) in the integrated genomic map. Further, since QTL that were coarsely mapped to vast stretches of the genome may mask any potential trends in genomic QTL distribution, QTL with total interval lengths that were statistical outliers were excluded according to the outlier formula: Upper limit = Upper quartile + (1.5*IQR), where the upper quartile is the 75^th^ percentile of QTL lengths and the IQR is the interquartile range of QTL lengths (75^th^ percentile – 25^th^ percentile). Next, the 25 chromosomes of the AstMex3 surface genome were divided into 118 non-overlapping 10 Mb windows using the tileGenome function from the GenomicRanges R package (v. 1.49.0; Lawrence et al. 2013). A window size of 10 Mb was chosen because it approximates the typical length of a QTL interval in the filtered dataset (mean length = 11.4 Mb, median length = 9.64 Mb). Windows were “centered” by finding the largest whole number of 10 Mb windows that could fit in each chromosome, subtracting the total length of these windows from the total length of the chromosome to find the “remaining” chromosome length, and shifting the starting position of the first window to a value equal to one half of the remaining length.

The number of QTL overlapping each window was then identified using the bed_intersect tool in the R package valr (v. 0.6.6; Riemondy et al. 2017). For the purposes of counting QTL within each window, all QTL for phenotypes relating to eye size (e.g., eye size, lens size, pupil area, etc.) were counted as one “distinct” QTL, as were all QTL for a melanophore phenotype (e.g., eye melanophore, dorsal melanophore, anal melanophore, etc.). The number of QTL overlapping each window after adjusting for these similar phenotypic traits is hereafter referred to as the number of “distinct QTL”.

To test whether the number of observed distinct QTL in each window matches a distribution expected by chance, a Monte-Carlo Chi-square test was conducted with the xmonte function from the XNomial R package (v 1.0.4; Engels 2015). Two expected distributions were tested: (1) a null expectation that QTL are distributed uniformly across the genome, and (2) the expectation that QTL are distributed across the genome proportional to protein-coding gene density.

Following the tests, 10 Mb windows with high standardized residual values (>3) against either the uniform distribution or the gene density distribution were grouped together and designated as putative QTL clusters (Table S7) (as in Peichel and Marques 2017). The observed distribution of distinct QTL across the *A. mexicanus* genome relative to the expected number of QTL under each of the two null expectations was visualized using the karyoploteR package (v. 1.24.0; Gel and Serra 2017).

### Testing potential molecular factors underlying QTL distribution

Mutational biases or opportunities may predispose regions to contribute to phenotypic evolution (Xie et al. 2019; Storz et al. 2019; Wortel et al. 2023). To investigate whether any molecular factors may explain the distribution of cave-associated QTL across the *A. mexicanus* genome, data was generated for each of the 118 10Mb windows of the AstMex3 surface genome assembly (Table S8). For each factor listed below, we conducted a Wilcox test in R to test for differences between the set of 10 Mb windows that were designated as putative QTL clusters and the set of 10 Mb windows that contain no mapped QTL. Six 10 Mb windows across four different chromosomes were identified as putative QTL clusters due to having significantly more QTL than expected under either the uniform null distribution or the gene density null distribution.

The following variables were investigated:

- *Repetitive DNA*: Repetitive DNA was quantified for each window as the proportion of all bases in the window that were reported as an “N” base in the hard-masked version of the AstMex3 surface fish genome, which was downloaded from the Ensembl Rapid Release FTP site (Cunningham et al. 2022). This method quantifies repetitive DNA in a broad sense, as it captures any sequence considered “repetitive” in the original masking procedures of the AstMex3 genome assembly.
- *GC content*: GC content was calculated for each window as the proportion of all bases in the window that are a “C” or “G” in the unmasked version of the AstMex3 surface fish genome.
- *CpG sites*: CpG sites were quantified for each window as the proportion of all dinucleotides in the window that are “CG” in the unmasked genome. Dinucleotide distributions were calculated using the oligonucleotideFrequency function from the Biostrings R package (v 2.66.0; Pages et al. 2016).
- *TGTGTG repeats*: Similarly, TGTGTG repeats are linked to replication fragile sites (Xie et al. 2019) and were quantified for each window as the proportion of all oligonucleotides of length 6 that are “TGTGTG” or “CACACA” in the unmasked genome with the oligonucleotideFrequency function.
- *Gene number*: The total number of genes, both protein-coding and non-protein-coding, overlapping each window was obtained with the bed_intersect function in the valr package (Riemondy et al. 2017), with the genome annotation file from AstMex3 (Ensembl rapid release, GTF format; (Cunningham et al. 2022)) providing the genomic coordinates of each gene. Gene counts were also obtained separately for genes of the following biotypes as reported in the GTF file: protein-coding, miRNA, snRNA, lncRNA.
- *Gene length*: The average and median protein-coding gene length in bp was calculated for each window using the start and end points given in the GTF file (i.e., the lengths included the entire length of the gene, not just the coding regions).
- *Transcription factor gene number*: Changes in the coding sequences of transcription factors (TF) may be involved in the evolution of gene regulatory networks, potentially facilitating phenotypic change (Cheatle Jarvela and Hinman 2015). The list of all genes annotated as transcription factors in *A. mexicanus* was downloaded from the Animal TFDB 4.0 (Shen et al. 2023). The bed_intersect function was then used to identify the number of TF genes that overlap with each window.
- *Recombination rate*: Linkage maps were generated for Pachón x Surface, Tinaja x Surface, and Molino x Surface F2 crosses as part of a separate ongoing study. For each linkage map, a recombination rate between every consecutive pair of markers was estimated by dividing the distance between the markers in centimorgans (cM) by the distance in Mb, giving a recombination rate in units of cM/Mb. These values were then used to find the average recombination rate across each 10 Mb window using a weighted average, where the weight applied to each individual rate was equal to the number of bp by which the interval overlapped the 10 Mb window.

### Analyzing effect sizes of cave-derived QTL

The distributions of reported PVE values for all QTL, as well as QTL in the Eye, Pigment, Skeletal, Non-Eye Sensory, Metabolic, and Behavior categories separately, were visualized as a histogram using ggplot2. These distributions were tested for skewness (a measure of the symmetry of a distribution) and kurtosis (a measure of whether the distribution is light-tailed or heavy-tailed) using the respective functions in the R package moments (v. 0.14.1; Komsta and Novomestky 2015).

### Identifying candidate genes underlying repeated cave-derived trait evolution

Lastly, we demonstrate the utility of our compiled QTL analysis for identifying candidate genes for future functional analysis. By cross-referencing QTL with other sources of genomic data, we identified candidate genes for cave-derived phenotypes in cavefish. First, the set of all genes overlapping with a QTL was obtained using the valr package in R (v. 0.6.6; Riemondy et al. 2017). Second, the set of all genes exhibiting evidence of selective sweeps in three independent cave populations (Pachón, Tinaja, and Molino) yet no evidence of selection in two surface populations (i.e., Río Choy, Rascón) was defined as in Moran et al. (2023). The Ensembl gene IDs published by Moran et al. (2023) were given for version 2 of the *A. mexicanus* genome. These IDs were converted to the version 3 equivalents using the AstMex3 homology table available on the Ensembl Rapid Release FTP site. Third, RNAseq data, originally collected by (Mack et al. 2021) from whole body samples of larval Pachón, Tinaja, and Molino cavefish as well as Río Choy surface fish, were reanalyzed. Raw reads were downloaded from the NCBI Sequence Read Archive (accession numbers given in Table S9 (Leinonen et al. 2010)), aligned to the AstMex3 surface fish genome using STAR (v. 2.7.1; Dobin et al. 2013), and aligned reads were counted using HTSeq (v. 0.11.0; Anders et al. 2015). Read counts were analyzed for differential expression between Pachón and Río Choy, Tinaja and Río Choy, and Molino and Río Choy, all at time 0 in the original study, using the DESeq2 package in R (v. 1.38.1; Love et al. 2014) (Tables S10, S11, S12), and differentially expressed genes were identified using a cutoff of a Benjamini-Hochberg adjusted p-value < 0.1. The set of genes that were differentially expressed in all three cave populations relative to the surface population, and for which the change in expression followed the same pattern in each population (i.e., upregulated in all three caves or downregulated in all three caves), were designated as genes with shared differential expression. Candidate genes for cave-derived phenotypes (Table S13) were identified by finding the subset of genes that 1) Have undergone shared selective sweeps in three separate cave populations and not in surface populations as defined by Moran et al. (2023), 2) Have shared differential expression between three cave populations relative to a surface population, and 3) Are found within a QTL for cave-derived traits.

## Results

### A map of cave-derived QTL across the Astyanax mexicanus genome

The first goal of this study was to provide a comprehensive map of previously published QTL for cave-derived traits in *A. mexicanus* to a new high-quality genome assembly, AstMex3, to facilitate analyses of trait evolution in this species. To this end, a total of 9023 genetic marker sequences from prior QTL studies (Tables 1, S1, S2) were collected and aligned to the AstMex3 surface fish genome using BLASTn (Table S3). Two-hundred markers did not have any BLAST hits, 126 markers had a BLAST hit that mapped to a chromosome that was discordant with other markers on the original linkage group, and 96 markers did not have a best BLAST hit that could be unambiguously resolved. The final integrated genomic map consisted of the best BLAST hit for the remaining 8486 genetic markers (Table S4).

This genomic map was used to associate 206 previously published QTL for cave-derived traits to the AstMex3 surface genome (Tables S4, S5). For 168 of the QTL, genetic markers marking the ends of the LOD confidence interval were part of the integrated genomic map, and the genomic range spanning the beginning of the first marker to the end of the second marker was denoted as a QTL interval. For the remaining 38 QTL, only a genetic marker associated with the peak of the LOD curve was present in the integrated genomic map, and the genomic range of this individual marker was denoted as a QTL peak. Table S6 summarizes the phenotypic category of each QTL that was mapped to the AstMex3 genome in this analysis. The resulting QTL map shows a general tendency for QTL for cave-derived traits to overlap with each other across the *A. mexicanus* genome (Figure S1, Tables S4, S7).

### Cave-derived QTL cluster in the Astyanax mexicanus genome

To test whether cave-derived QTL cluster in the *A. mexicanus* genome more than expected by chance, the observed QTL distribution across 10 Mb genomic windows was tested for fit against two expected distributions. First, we found that the observed QTL distribution does not fit the expected distribution under a null model which assumes a uniform distribution of QTL across the genome (chi-square = 192.9, p = 0.00002; Figure 1). Second, the observed QTL distribution does not fit the expected distribution under a model in which QTL density is proportional to protein-coding gene density (chi-square = 216.1, p < 0.00001; Figure 1). Together, these results support the hypothesis that QTL for cave-derived traits cluster in the genome. Six 10 Mb windows across four different chromosomes were identified as putative QTL clusters due to having significantly more QTL than expected under either distribution (Table S7).

**Figure 1.**
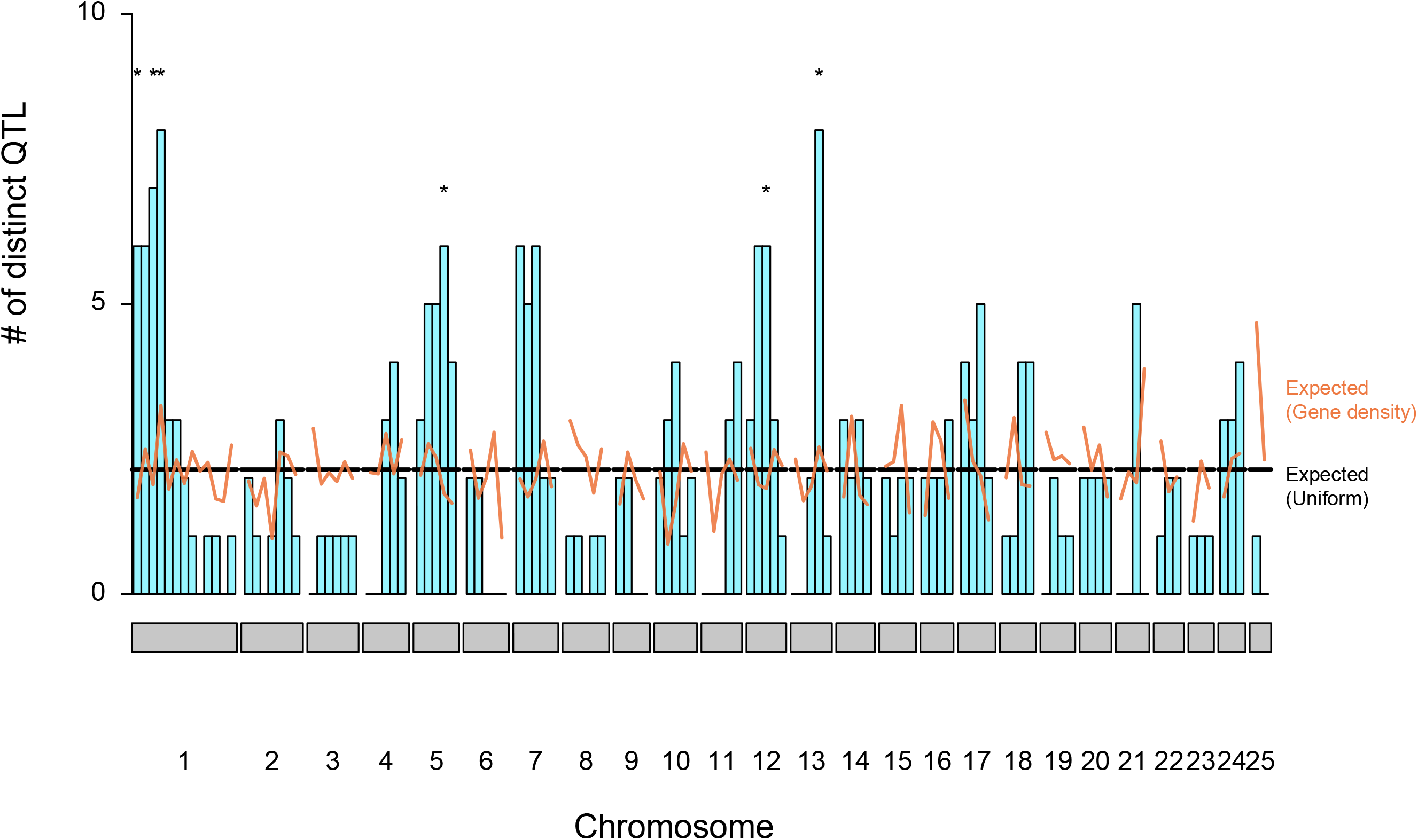
Observed distribution of the number of distinct cave-associated QTL in each 10 Mb window of the *Astyanax mexicanus* genome (cyan histogram) compared to the expected distribution under a null model in which the QTL distribution is uniform across the genome (black line), and a model in which the QTL distribution is proportional to protein-coding gene density (orange line). Asterisks denote specific 10 Mb windows which contain more QTL than expected under either of the models (standardized residual > 3).

### Distribution of putative QTL clusters is not explained by molecular factors

Increased mutation rates lead to increased opportunity for evolution (Kimura and Ohta 1971; Bromham 2009). To investigate whether the distribution of putative QTL clusters across the *A. mexicanus* genome are associated with known characteristics of the genome that are potentially associated with increased opportunity for mutation, 13 variables including recombination rate, gene length, gene number, transcription factor number, CpG sites, TGTGTG repeats, and repetitive DNA were tested for significant differences between putative QTL clusters (n=6) compared to 10 Mb windows that contain no known QTL (n=22; Table S8). None of the variables tested were significantly different between the two groups (Wilcoxon rank-sums test, p>0.05 for all tests), suggesting that none of these factors are associated with cave-derived evolutionary “hotspots” at this coarse scale (Figure S2).

### Distribution of effect sizes depends on type of trait

While acknowledging that we will miss many small and medium effect loci, the distribution of percent variance explained (PVE) values reported for all previously mapped cave-derived QTL in *A. mexicanus* was compiled to understand more about the distributions of effect size across different trait classifications (Figure 2). The distribution is extremely positively skewed (Skewness coefficient = 2.04) and light-tailed (kurtosis coefficient = 8.02). This indicates that most alleles detected to be associated with cave-derived phenotypes explain a relatively small proportion of phenotypic variation, with just a few outlying mutations that contribute more substantially to cave-derived phenotypes.

**Figure 2.**
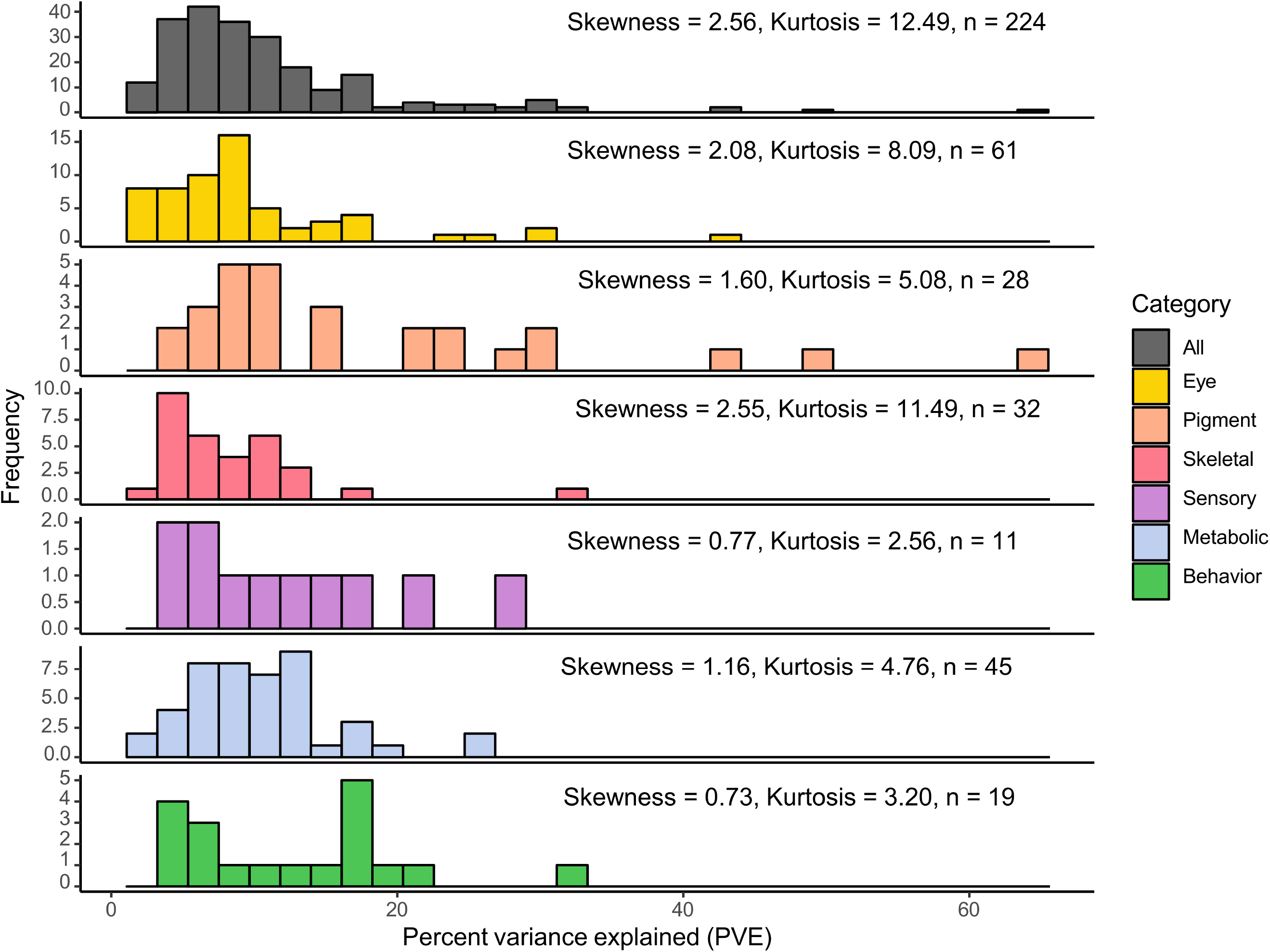
Distribution of reported percent variance explained (PVE) values for previously published cave-associated QTL in *Astyanax mexicanus* by trait category. Skewness coefficients indicate the degree of asymmetry in the distribution; in this case, all distributions are positively skewed, indicating that there are more mutations of below average effect size than of above average effect size. Kurtosis coefficients indicate the degree to which the distribution is light-tailed relative to a normal distribution, (i.e., higher values of kurtosis mean that outlying mutations with high PVE are more frequent).

This trend generally holds true when cave-derived QTL are divided into trait categories – all distributions are positively skewed and light-tailed, but there is variation in the degrees of skewness and kurtosis. The most highly positive skewed trait categories are Eye (skewness coefficient = 2.08) and Skeletal (skewness coefficient = 2.55), suggesting that a strong majority of mutations affecting these traits are of small effect with a few mutations having larger effects. The least strongly positively skewed trait categories are Non-Eye Sensory (skewness coefficient = 0.77) and Behavior (skewness coefficient = 0.73), suggesting that the distribution of effect sizes for these trait categories are somewhat larger than other traits. Notably, traits in the Pigment category have some of the largest effect sizes across all traits (Figure 2), and this may be because multiple studies mapped albinism which is a Mendelian trait.

### A robust list of candidate genes underlying repeated cave-derived trait evolution

By utilizing multiple avenues of genomic data, the pool of candidate genes underlying traits of interest can be narrowed. We identified 36 candidate genes (Table S13) underlying the repeated evolution of cave-derived traits in *A. mexicanus* by finding the subset of protein-coding genes that are present in at least one cave-derived QTL, have evidence of repeated selective sweeps across three populations of cavefish (Moran et al. 2023), and are differentially expressed between cavefish and surface fish in all of the same three cave populations (Mack et al. 2021) (Tables S10-S12).

## Discussion

The genetic architecture underlying adaptive evolution is a central topic in evolutionary biology. QTL studies can identify regions of the genome important for trait evolution and, thus, are a powerful method for investigating genetic mechanisms of adaptation. In this study, 206 quantitative trait loci (QTL) from 16 previously published studies (Protas et al. 2008; Protas et al. 2006; Protas et al. 2007; Gross et al. 2009; Yoshizawa et al. 2012; O’Quin et al. 2015; O’Quin et al. 2013; Kowalko et al. 2013a; Kowalko et al. 2013b; Warren et al. 2021; Stockdale et al. 2018; Carlson et al. 2018; Yoshizawa et al. 2015; Powers et al. 2020; Gross et al. 2014; Lyon et al. 2017) were mapped to the newly published *A. mexicanus* surface genome assembly, AstMex3. Collectively, these QTL represent a variety of morphological, physiological, and behavioral traits of interest that differentiate Mexican cavefish from their surface-dwelling counterparts. Our study organizes the disparate maps onto one high quality genome assembly, allowing for easy comparison and analysis of QTL features across studies. Further, this work is an improvement on a previous QTL review in *A. mexicanus* because it maps QTL to a more contiguous surface fish genome, capturing loci in QTL intervals that may have been fragmented or may not have been present in the cavefish genome (O’Quin and McGaugh 2015).

This integrated QTL map showed that the observed distribution of QTL for cave-derived traits across the *A. mexicanus* genome is non-uniform and is not proportional to protein-coding gene density. As a result of this analysis, six putative QTL clusters that contain more unique QTL than expected by chance were identified on four of the 25 chromosomes in the *A. mexicanus* genome. The most striking of these clusters occurs on Chromosome 1 from base pairs ∼2Mb-42Mb. This 40 Mb region contains QTL for 10 distinct phenotypic traits including eye size, vibration attraction behavior, relative condition, tastebuds, melanophore, and several others, suggesting that Chromosome 1 is a hotspot of cave trait evolution in *A. mexicanus*. This result is very similar to the conclusion of a similar analysis in three-spined stickleback, which found that five out of 21 chromosomes contained more QTL than expected by chance (Peichel and Marques 2017), suggesting that genomic hotspots drive adaptation in both of these systems.

Clustering of QTL for seemingly distinct traits, such as what is observed on Chromosome 1, could be due to pleiotropic loci that influence multiple phenotypic traits of interest (O’Quin and McGaugh 2015; Peichel and Marques 2017). Pleiotropy could help explain adaptation to the cave environment that has independently occurred in multiple populations of this species, as selective pressure on one trait could drive coordinated change in other traits and bring about extensive phenotypic change (Yamamoto et al. 2009; Jeffery 2005). The putative QTL clusters identified in this study provide an ideal starting point to search for genes that may have pleiotropic effects (Jeffery 2010; Yoshizawa et al. 2013; Protas et al. 2007). One such pleiotropic gene, *oca2*, has already been identified in the literature and is known to affect both albinism and sleep duration (O’Gorman et al. 2021). Alternatively, tightly linked genes may underlie the traits implicated in QTL clusters (Borowsky 2013), and indeed many suites of traits in other systems have been found to be in tight physical linkage or genomic inversions (Noor et al. 2001; Wellenreuther and Bernatchez 2018; Wilder et al. 2020). Functional interrogation, for example with CRISPR-Cas9 gene editing (Klaassen et al. 2018) of genes contained in QTL clusters, represents a promising next step toward differentiating between these two hypotheses.

More broadly, the integrated QTL map is useful in analyzing how the distribution of cave-derived QTL correlates with other factors on a genomic scale. Certain molecular attributes may influence which regions of the genome are more likely to mutate and contribute to phenotypic evolution, including recombination rate (Lercher and Hurst 2002; Halldorsson et al. 2019) abundance of CpG sites (Storz et al. 2019), TGTGTG repeats (Xie et al. 2019), and gene length (Woodhouse and Hufford 2019; Moran et al. 2023; Mei et al. 2018). If supported, these hypotheses suggest mutational biases predispose certain regions of the genome to phenotypic evolution. In our study, however, none of the 13 variables that were tested were significantly correlated with the distribution of QTL on a genomic scale. While these correlates exhibit no significant correlations to QTL clusters, the corresponding QTL intervals in the integrated map presented here are quite large and overlap several hundred genes. Thus, only a subsection of the QTL region may contribute in a meaningful way to the phenotype and the scale at which we examined the correlations may be too broad to accurately address correlations between genomic features and cave-derived evolution.

The collection of all previously published QTL into one dataset facilitates the analysis of effect sizes of cave-derived QTL across a range of traits. While most adaptative walks occur with predominantly small effect mutations, large-effect mutations may be utilized when responding to large environmental shifts (Orr 2006), especially when, as in the cavefish system, migration with non-adapted populations occurs (Yeaman and Whitlock 2011). To assess whether large-effect mutations may play an outsized role in the evolution of certain cave-derived phenotypes, the percent variance explained (PVE) reported for each QTL was recorded for all studies identified in the literature review. The QTL with the highest PVE value is a QTL for albinism, which is a Mendelian trait attributed to *oca2* (Protas et al. 2006), and pigmentation related traits seem to have several large-effect outliers compared to other trait classes. The phenomenon of a single gene explaining a large amount of variation in a trait, like *oca2* does for albinism, is the exception, similar to the QTL review of stickleback fish (Peichel and Marques 2017).

Additionally, QTL mapping experiments tend to overestimate PVE (Beavis 1994) and be biased in detecting mostly larger-effect alleles (Rockman 2012), therefore, some of the outlying QTL with high PVE values may not have a true effect size as great as their position in the distribution suggests. A final caveat is that the bias in overestimating PVE values increases with decreasing sample size, so comparing PVE effects of studies with different sample sizes is inexact (Beavis 1994; Xu 2003). Despite the bias in detecting large effect alleles, the shape of the distribution of PVE values across several traits and studies suggests that adaptation is generally attributable to many mutations of small effect rather than few mutations of large effect in *A. mexicanus*.

Likely most important for evolutionary geneticists, the integrated QTL map presented in this study provides an ideal starting point for identifying candidate genes underlying cave-derived traits. As proof of principle, here we utilized three sources of data – QTL, differential expression, and population genomics – to curate a list of just 36 genes that are likely to be involved in the repeated evolution of cave-derived traits in *A. mexicanus* (Table S13). Probably the most exciting candidate is *rgrb*, which encodes retinal G protein-coupled receptor b predicted to be an opsin and is widely expressed across zebrafish tissues (Davies et al. 2015). We find that *rgrb* maps to a QTL for retinal thickness (specifically the outer plexiform layer (OPL)), and that *rgrb* is downregulated in cavefish populations relative to surface populations in whole body samples from 30dpf fry. To further investigate gene expression trends in *rgrb*, we re-mapped an RNAseq dataset collected from eye tissue of developing embryos at 54 hpf to the AstMex3 genome assembly ((Gore et al. 2018); see methods for RNAseq re-mapping procedure, Table S14), and we confirmed that *rgrb* is downregulated in developing eye tissue of Pachón cavefish relative to surface fish. As with all candidates on our list, *rgrb* exhibits evidence of selective sweeps across three cave populations, suggesting that *rgrb* is a target of adaptation to the cave environment.

Interestingly, exon 4 and exon 5 are nearly completely deleted across multiple cave populations (Lineage 1: Escondido, Molino, Jineo; Lineage 2: Jos, Monticellos, Pachón, Palma Seca, Tinaja, Yerbaniz), while reading frames are intact for all surface populations (see supplementary alignment). Together, these lines of evidence strongly suggest that *rgrb* has been a target of repeated selection in *A. mexicanus*, and that exon deletions and downregulation of *rgrb* may be involved in convergent abnormal retinal development in *A. mexicanus* cavefish (Emam et al. 2020).

Other intriguing candidate genes from our list span a variety of traits. For instance, we find that the retinoic acid receptor *stra6* is downregulated in cavefish and falls under a QTL for maxillary bone length, which tends to be longer in cavefish than surface fish (Atukorala et al. 2013). Retinoic acid signaling is known to play a role in skull and tooth development in fish (Draut et al. 2019), so it is possible that *stra6* downregulation may influence maxillary bone morphology in *A. mexicanus* through this avenue. As another example, the rab-GTPase *rab8a* has been implicated in the maintenance of glucose homeostasis through its regulation of glucose transporters (Sun et al. 2010), which could connect to our finding that *rab8a* is upregulated in cavefish and is under a QTL for relative condition, a measure of observed vs predicted weight in which cavefish tend to be higher than surface fish (Protas et al. 2008). *Rab8a* is also associated with a QTL for peduncle depth. Many of the candidate genes on our list are associated with QTL for more than one phenotype, raising the possibility of pleiotropic effects like what has been observed in *oca2* (O’Gorman et al. 2021). While we have highlighted just a few examples, they demonstrate the utility of the data presented here in guiding future functional characterization.

In summary, the results presented in this study suggest that adaptation to the cave environment in populations of *A. mexicanus* has been driven by alleles in concentrated regions of the genome. While there are no molecular factors that reliably predict the hotspots of cave-derived trait evolution on a genomic scale, candidate genes for repeated trait evolution represent a promising avenue to study repeated targets in adaptation to extreme environments. These results contribute to the goal of understanding adaptation to extreme environments on a genomic scale and will serve as a useful resource to guide future studies on the genetic basis of cave-derived trait evolution in *A. mexicanus*.

## Data availability

All data and analyses are provided in the supplemental materials.

## Acknowledgements

We thank the Minnesota Supercomputing Institute without which this work would not be possible. We thank Jeff Gralnick, Yaniv Brandvain, and the McGaugh lab for comments on earlier drafts of this manuscript.

